# Examining Changes in Gramicidin Current Induced by Endocannabinoids

**DOI:** 10.1101/2024.11.05.622072

**Authors:** Sultan Mayar, Audrey Cyr-Athis, Nazzareno D’Avanzo

**Author notes:** **Correspondence:** Nazzareno D’Avanzo.

## Abstract

Endocannabinoids are a diverse family of lipid molecules, which circulate in the human body, impacting the cardiovascular and the nervous systems. Endocannabinoids can influence pain perception, appetite, stress responses, mood, memory and learning. Regulation of these lipids present a promising therapeutic avenue for numerous neurological disorders. In addition to acting as agonists to cannabinoid receptors (CBRs), endocannabinoids can also modulate the function of various ion channels and receptors independently of CBRs. This modulation of function can arise from direct binding to the channel proteins, or via changes to the lipid properties such as membrane elasticity/stiffness, curvature, or hydrophobic thickness. Here, we assess the effects of endocannabinoids on membrane properties by examining changes in gramicidin (gA) currents in *Xenopus* oocytes. Endocannabinoids from both classes (Fatty acid ethanolamides (FAEs) and 2-monoacylglycerols (2-MGs)) are studied and current-voltage relationships are assessed. Employing gramicidin channels as molecular force probes can enable both predictive and quantitative studies on the impact of bilayer-mediated regulation on membrane protein function by endocannabinoids.

## INTRODUCTION

Endocannabinoids are lipid molecules which are naturally synthesized in the human brain and peripheral tissues (Ruiz de Azua and Lutz 2019). Two other cannabinoid classes are exogenous or phytocannabinoids, which are extracted from plants, and synthetic cannabinoids, which are generated industrially as potent agonists. Being a part of the endocannabinoid system (ECS), endocannabinoids are responsible for a variety of different mechanisms in regulating human body homeostasis. These lipid-like molecules have been linked to modulating metabolism, obesity, appetite, stress responses, mood, memory and learning and neurodegenerative disorders (Romano et al. 2014; Rahman et al. 2021; Cristino, Bisogno, and Di Marzo 2020). Endocannabinoids are directly derived from membrane phospholipids (phosphatidylinositol 4,5-bisphosphate, PIP_2_ and phosphatidylethanolamine, PE) and are known to modulate cell signalling via cannabinoid receptors (CBRs) (Howlett 2002; Ruiz de Azua and Lutz 2019). Mounting evidence has also pointed to endocannabinoids modulating ion channels and receptors independently of CBRs. For example, various endocannabinoids have demonstrated effects in TRP channels (Muller, Morales, and Reggio 2019; Zygmunt et al. 1999), Kv channels (Iannotti et al. 2014; Oliver et al. 2004), Cav channels (Shimasue et al. 1996; Chemin et al. 2001) and more recently Kir channels (Mayar et al. 2024) expressed in heterologous systems that do not express CBRs.

Many studies of cannabinoid regulation of channel function suggest effects via direct interactions (via binding site) or indirect (via signalling cascade) mechanisms. However, in addition to signalling pathways, cannabinoids may also act by altering the physiochemical properties of the lipid membrane (Oz, Yang, and Mahgoub 2022). For instance, the exogenous cannabinoid, cannabidiol (CBD) which is the nonpsychoactive component of marijuana (ElSohly 2007), has been shown to modulate Nav1.4 channels via both direct interactions (i.e. pore binding) and via changes in membrane properties, which then alters fenestrations in the bilayer-spanning domain of the channel (Ghovanloo et al. 2021; Ghovanloo, Goodchild, and Ruben 2022). A recent examination of endocannabinoid regulation of Kir channels using surface plasmon resonance and molecular dynamics simulations demonstrates these molecules do not directly interact with these channels (Mayar et al. 2024). Therefore, it is possible that endocannabinoids may also exert their effects through changes in membrane properties.

Monitoring changes in gramicidin currents (gA) across different membrane potentials has been demonstrated to be an effective tool in assessing the ability for compounds to modulate membrane properties (Lundbæk et al. 2004; Kapoor et al. 2019; Ghovanloo, Goodchild, and Ruben 2022). Gramicidin which was originally identified as an antibiotic (Dubos 1939; Kelkar and Chattopadhyay 2007) allows for the permeation of water and cations through a ∼4 Å functional channel pore which spans the lipid membrane. To create this channel, two gramicidin monomers (one in each side of membrane leaflet) need to dimerize (Fig. 1) (Sawyer, Koeppe, and Andersen 1989; Lundbæk et al. 2004). Preference for dimerization or dissociation back to individual monomers can be directly correlated to membrane properties such elasticity or stiffness (Andersen and Koeppe 2007; Sawyer, Koeppe, and Andersen 1989). Therefore, changes in gramicidin dimerization can be directly related to changes in membrane properties. Notably, amphiphilic molecules which share both lipophilic and hydrophilic properties such as Triton X 100 (TX-100) have been shown to modulate membrane properties and affect gramicidin currents (Sawyer, Koeppe, and Andersen 1989; Lundbæk et al. 2004; Ashrafuzzaman, Koeppe, and Andersen 2024). Endocannabinoids are also considered amphiphilic entities which have a carbon tail and a polar head group (Basavarajappa 2007).

**Figure 1:**
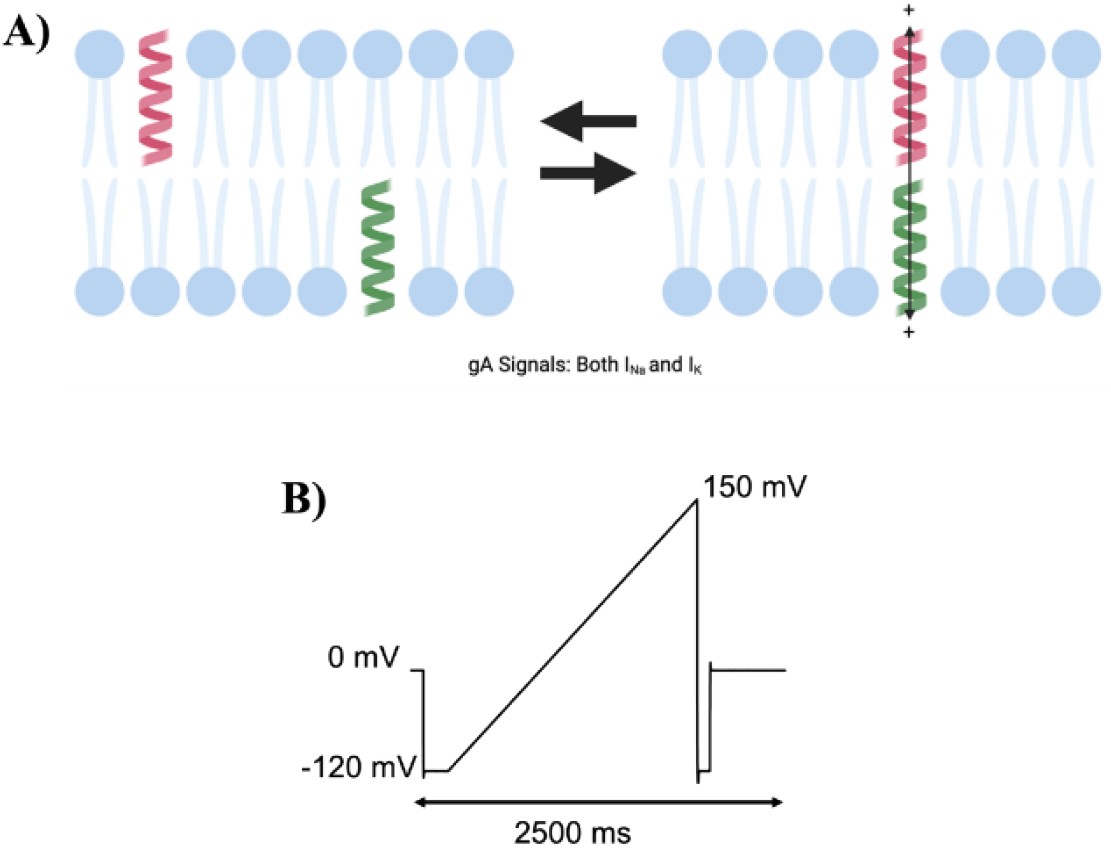
Gramicidin (gA) currents used to monitor changes in membrane elasticity/stiffness. **A)** Cartoon of gramicidin dimerization to form a conducting cationic pore. **B)** Voltage protocol applied to obtain gramicidin current. Cells were held at 0 mV, followed by a step down to -120 mV and ramping up to +150 mV over 2500 milliseconds. gA currents were assessed before and after the addition of endocannabinoids in pair-wised experiments.

In this observational study, we examine whether two classes of endocannabinoids, 2-monoacylglycerols (2-MGs) and fatty acid ethanolamides (FAEs) modulate membrane properties by assessing changes macroscopic gramicidin (gA) current. It has been previously demonstrated that amphiphiles such as TX100 effect membrane elasticity, largely through changes in the bilayer spring constant (Lundbæk et al. 2004; Lundbæk, Koeppe, and Andersen 2010). Using a *Xenopus* oocyte cell-model and two-electrode voltage clamp (TEVC) we elicited cationic gramicidin currents in the presence and absence of various endocannabinoids and correlated changes with the structural properties of these lipid-like molecules.

## MATERIALS & METHODS

### Drugs and reagents

Solid gramicidin was dissolved in extracellular recording solution to a final concentration of 65 µM and was made fresh on the day of experiments (Sigma-Aldrich, USA). All endocannabinoids were pre-diluted in 99.8% DMSO at a varying concentration (Cayman Chemicals, USA). The final concentration of each endocannabinoid tested was 30 µM in the bath. Detergent, Triton™ X-100 (Sigma-Aldrich, USA) was diluted to a working concentration of 10 mM with distilled water to form a stock solution. The final concentration of Triton™ X-100 added to the bath was 20 µM, which has been previously demonstrated to alter membrane properties (Sawyer, Koeppe, and Andersen 1989; Lundbaek et al. 1996; Lundbæk et al. 2004).

### Oocyte Preparation

Experiments were performed using unfertilized *Xenopus* oocytes, as these have been demonstrated to lack cannabinoid receptors (Karimi et al. 2018; Peshkin et al. 2019). Cells were surgically extracted from female *Xenopus laevis* frogs which were anesthetized with tricaine methanesulfonate (Sigma-Aldrich, USA). After extraction of oocyte sacks, cells were treated with collagenase type IA (Sigma-Aldrich, USA) Stage IV and V oocytes were manually sorted and placed in a vial containing Barth antibiotic solution supplemented with 5% horse serum (mM): 90 NaCl, 3 KCl, 0.82 MgSO_4_.7H_2_O, 0.41 CaCl_2_.2H_2_O, 0.33 Ca(NO_3_)_2_.4H_2_O and 5 HEPES supplemented with 100 U/mL of penicillin-streptomycin and 10 mg/mL of kanamycin stock (10 mg/mL). Oocytes were placed in a control temperature of 17 – 19 °C prior to experiments.

### Electrophysiological Recordings

Uninjected *Xenopus* oocytes were for two-electrode voltage clamp (TEVC). Glass borosilicate rapid fill microelectrode pipettes containing a 1 M KCL solution were used to impale cells and measure gramicidin (gA) currents. Briefly, during recordings oocytes were placed in a bath containing an external recording solution (in mM): 5 KCl, 84 NaCl 15 HEPES, 0.4 CaCl_2_, and 0.8 MgCl_2_, pH = 7.4. Gramicidin currents were elicited using a 2.5 second repetitive pulse protocol holding our cells at 0 mV, stepping down to -120 mV and ramping to +150 mV. Inter-pulse time for each pulse was 30 seconds to allow for the endogenous membrane and proteins to fully recover after each sweep. Currents were amplified by an OC-725C amplifier (Warner Instruments, USA) and digitized using a Digidata 1322A (Molecular Devices). Data were obtained with Clampex 10.5 at a sampling rate of 5 KHz with a filter of 1 KHz.

During electrophysiological recordings, the repetitive pulse protocol was taken as a control, followed by the addition of 65 μM of pre-diluted gramicidin. Additional pulse protocols were recorded in the presence of gramicidin and once the current density stabilized (typically 5-10 mins) the endocannabinoid was added into the extracellular bath solution at a final concentration of 30 μM. Recordings were completed once the current density stabilized following the addition of the endocannabinoid (typically after 30 minutes). Each recording was conducted at ambient room temperature.

### Data analysis and statistics

Raw cationic current recordings were analyzed offline using the Clampfit Software (Molecular Devices). Graphing was conducted using the Origin 9.0 Software (Northampton, MA, USA). Currents were measured at specified voltages (−120 mV, -100 mV, -50 mV, 0 mV, +50 mV, +100 mV and +150 mV) for each cell in the absence then presence of the specific endocannabinoid after current stabilization at each step. Post-treatment currents were normalized to the control current (before the addition of the endocannabinoid, 0 µM) at +150 mV. Normalized currents were then averaged for each endocannabinoid condition and plotted for I-V curved with standard error of means (SEM). This enables pair-wise analysis to determine whether the endocannabinoid effects gramicidin currents.

Endocannabinoids contain three structural characteristics: chain length (number of carbons), degree of unsaturation (number of double carbon bonds) and the position of the first unsaturated bond (Figure S1). To examine the correlation between the maximal change in gramicidin current at +150 mV versus these structural characteristics a linear relationship was utilized following the equation:

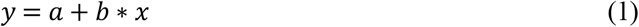

Where appropriate a non-linear Gaussian fit equation was used:

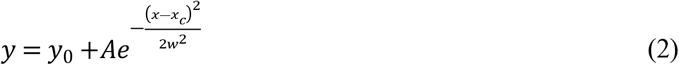

Changes in gramicidin current as a function of any 2 of the 3 characteristics were also plotted as three-dimensional (3-D) surface plots.

All data here are presented as means of (±) standard errors for the noted *n* recordings. Pairwise sample student t-test was used to determine statistical significance between the mean currents at each voltage for each test condition (endocannabinoid) versus its own control. The level of significance was denoted as α = 0.05. Any P-value less than 0.05 was denoted as statistically significant *, P < 0.01 denoted with ** and P < 0.001 denoted ***. Pearson correlation coefficient (R) and coefficient of determination (R^2^) are also reported for each linear fit.

## RESULTS

### Fatty Acid Ethanolamides (FAEs) endocannabinoids modulate gramicidin currents and membrane properties

In this study, we assessed changes in cationic currents through gramicidin channels induced by endocannabinoids by two-electrode voltage clamp of *Xenopus* oocytes. Unfertilized *Xenopus* oocytes do not express cannabinoid receptors (Karimi et al. 2018; Peshkin et al. 2019), and thus ensure that any changes we observe are due to changes in membrane properties, rather than via intracellular signaling induced by cannabinoid receptor activation. Gramicidin monomers dimerize to form conducting (sodium and potassium) cationic channels embedded in the membrane bilayer (Sun et al. 2019; Hladky and Haydon 1970) (Fig. 1). This dimerization can be directly correlated to membrane properties such as elasticity, stiffness and thickness (Ghovanloo, Goodchild, and Ruben 2022; Andersen and Koeppe 2007; Lundbæk et al. 2009). Oocytes were bathed in a physiological low potassium (5 mM) and high sodium (84 mM) external solution, and cationic currents were assessed using a repetitive ramp protocol started holding at 0 mV, stepping down to hyperpolarized potential of -120 mV and ramping to +150 mV (Fig. 1). Once these basal currents were stable, we then added 65 μM gramicidin to the extracellular bath and waited until gramicidin currents were stable (typically 5 mins) before adding 30 μM of the test endocannabinoid. Measurements were taken every 60 seconds and current stabilization typically required incubation for approximately 30 minutes. This time course was consistent with previous studies involving the addition of cannabinoids to *Xenopus* oocyte membranes (Mayar et al. 2024; Mayar et al. 2022). Currents before and after endocannabinoid treatment were compared in a pair-wise manner by normalizing the I-V for each cell to the maximal gramicidin current recorded at +150 mV prior to the addition of the endocannabinoid. These normalized I-Vs for each condition were then averaged and analyzed for statistical significance at each voltage. Amphiphiles such as Triton X-100, have been demonstrated to modulate membrane elasticity and alter gA current (Ghovanloo, Goodchild, and Ruben 2022; Sawyer, Koeppe, and Andersen 1989). TX-100 does not affect membrane thickness (Lundbæk, Koeppe, and Andersen 2010) nor unitary conductance of gA (Lundbæk et al. 2004). Rather, a decrease in gA current is associated with a decrease in membrane stiffness (conversely an increase in membrane elasticity) and an increase in gA current implies an decrease in membrane elasticity (or increase in membrane stiffness) (Sawyer, Koeppe, and Andersen 1990; Lundbaek et al. 1996; Lundbæk, Maer, and Andersen 1997; Sawyer, Koeppe, and Andersen 1989).

In oocytes, we observe a decrease in macroscopic gA currents following treatment with TX-100 (+100 mV: P = 0.003; +150 mV: P = <0.0001) (Fig. S1 and Table S1) suggesting that TX-100 alters membrane elasticity in oocytes thereby decreasing the number of conducting/functional dimers. Similar results were also observed in untransfected HEK cells (Ghovanloo, Goodchild, and Ruben 2022).

In this study, we examined the effects of endocannabinoids on gA currents as a surrogate for assessing membrane elasticity/stiffness. Endocannabinoids tested are organized in two chemical families which differ in their headgroup; fatty acid ethanolamides (FAEs) and 2-monoacylglycerols (2-MGs). Our findings reveal various FAE endocannabinoids modulate gA current with a diversity of responses. gA current traces before and after endocannabinoid treatment were normalized to current elicited at +150 mV in control (no endocannabinoid), and normalized current-voltage (I-V) relationships were determined (Fig. 2, Fig. 3 and Table S1). Interestingly, anandamide (AEA), which was the first and most studied endocannabinoid, show no change in gA current at any voltage (Fig. 2A and Table S1). Oxy-AEA, which is an oxyhomologue of anandamide, alongside NEA and LEA show significant increases in gA currents (∼20 - 40% at +150 mV; Fig. 2 and Table S1). α-LnEA, SEA and TEA had a more moderate effect on gA current, with increases of approximately 15% (Table S1). Interestingly, OEA is the only endocannabinoid which significantly reduced gA current by approximately 10% at +150 mV (P < 0.001), which indicates this endocannabinoid decreases dimerization and could increase membrane elasticity (Fig. 2D and Table S1). ArEA and γLnEA induce non-significant changes in gA current of ∼5% at +150 mV suggesting these endocannabinoids do not alter native oocyte membrane properties (Table S1).

**Figure 2.**
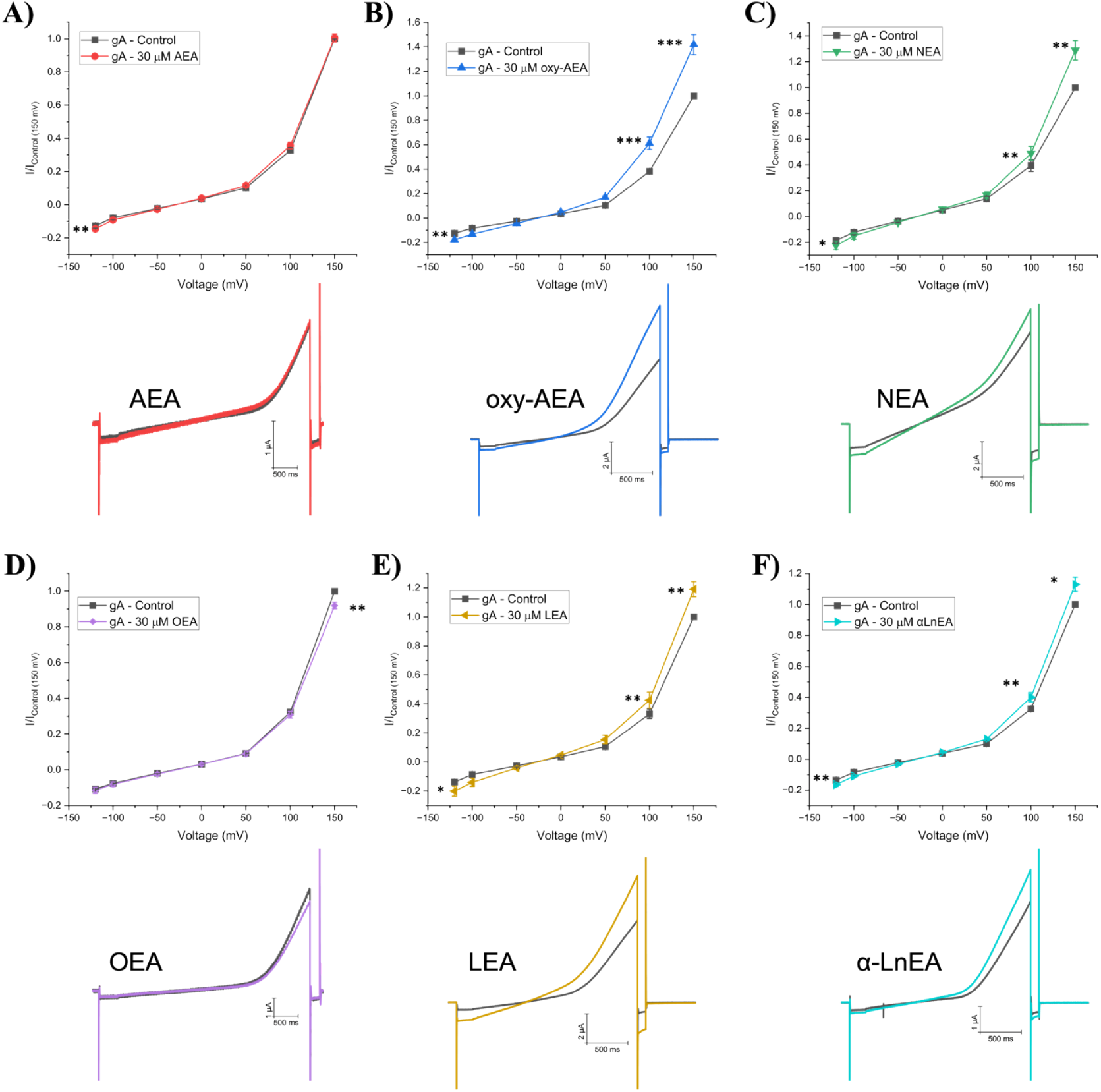
Fatty acid Ethanolamides (FAEs) effect on gramicidin currents (gA). (**A-F**) (*top*) Averaged cationic gramicidin currents before and after treatment with AEA, oxy-AEA, NEA, OEA, LEA or αLnEA respectively. Test currents were averaged at -120 mV, -100 mV, -50 mV, 0 mV, +50 mV, +100 mV, and +150 mV. (*below*) Representative gA current traces before and after treatment with AEA, oxy-AEA, NEA, OEA, LEA or αLnEA. Paired student t-test was used to determine I/I_max_ differences at -120, +100 and +150 mV (n > 6 < 12; * = P < 0.05, ** = P < 0.01, *** = P < 0.001)

**Figure 3.**
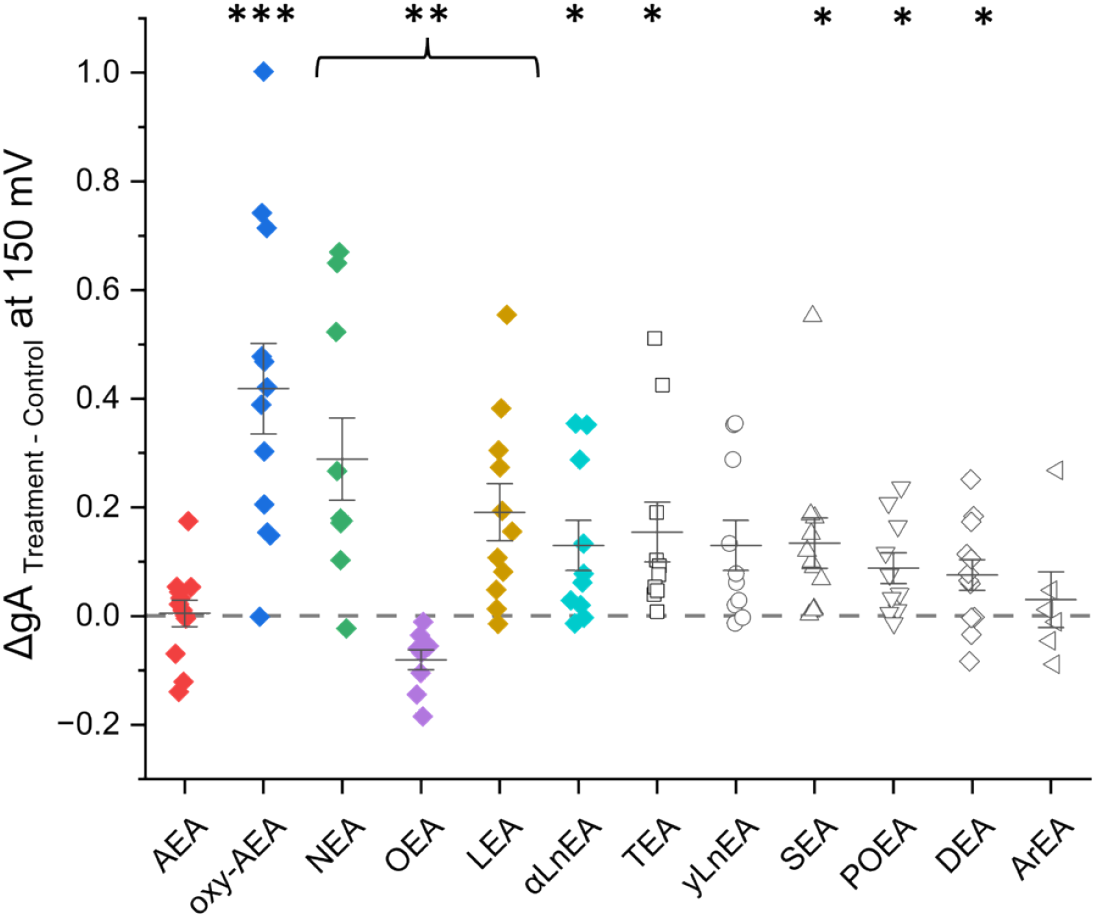
Effect of fatty acid ethanolamides (FAEs) on gramicidin currents (gA) at +150 mV. Difference in normalized cationic currents (endocannabinoid treatment vs control). All differences are plotted for +150 mV. Paired student t-test was used to determine gA differences between treatment and control (n > 6 < 12; * = P < 0.05, ** = P < 0.01, *** = P < 0.001)

Endocannabinoids can be characterized by their physiochemical including chain length, degree of unsaturation and the position of the first unsaturated bond (Fig. S2). We sought to determine whether the changes in gA current induced by FAEs correlate to any of these physiochemical properties (Fig. 3). Linear fits reveal no correlation between changes in gA current and either of the three properties (Fig. 3A-C, Table S3). This indicates that changes in membrane properties induced by FAEs are not driven by a single property. Therefore, we generated three-dimensional surface maps defining the relationship between gramicidin currents and pairs of the physiochemical properties of the FAEs (Fig. 3D-F). These profiles indicate that endocannabinoids with longer tails (20-24) and fewer unsaturated bonds (where unsaturation occurs far from the headgroup) lead to the greatest increase in gA current, and thus, decrease in membrane elasticity. The only FAE to decrease gA current (i.e. increase elasticity) is OEA which has an 18-carbon tail, 1 degree of unsaturation and the first carbon double bond at the 9^th^ position (Fig. S1).

### 2-monoacylglycerols (2-MGs) endocannabinoids modulate gramicidin currents and membrane properties

We also examined the effects of 2-monoacylglycerol endocannabinoids (2-MGs) on membrane properties. 2-MGs contain an glycerol as their headgroup typically bound at the *sn-2* position, though linkage at the *sn-1* position is possible as well. Average normalized I-V relationships are plotted alongside representative raw gA current traces in Fig. 4. Our data reveals varying effects of 2-MG endocannabinoids on gA current and thus, their ability to modulate membrane properties. Surprisingly, another well-characterized endocannabinoid, 2-arachidonoyl glycerol (2-AG) does not significantly change gramicidin currents at potential +100 mV (P = 0.19) nor +150 mV (P = 0.70) (Fig. 4A, D and Table S2). Therefore, 2-AG may not exert effects on other proteins through changes in membrane properties but rather through direct interactions with protein targets. 2-PG 2-SG, and 1-SG increase gA current and thus decrease membrane elasticity (P < 0.01) (Fig. 4 and Table. S2). 1-OrG is the only endocannabinoid in the 2-MG class which significantly decreases gramicidin currents (∼18% at +150 mV (P=0.02)), indicating a possible increase in membrane elasticity (Fig. 4B and Table S2). Five other endocannabinoids in this class (1-AG, 1-MrG, 1-OG, 2-LG and 2-SrG) do not alter gA current (Table S2) and thus do not alter the elasticity/stiffness of a the biological *Xenopus* oocyte membrane.

**Figure 4.**
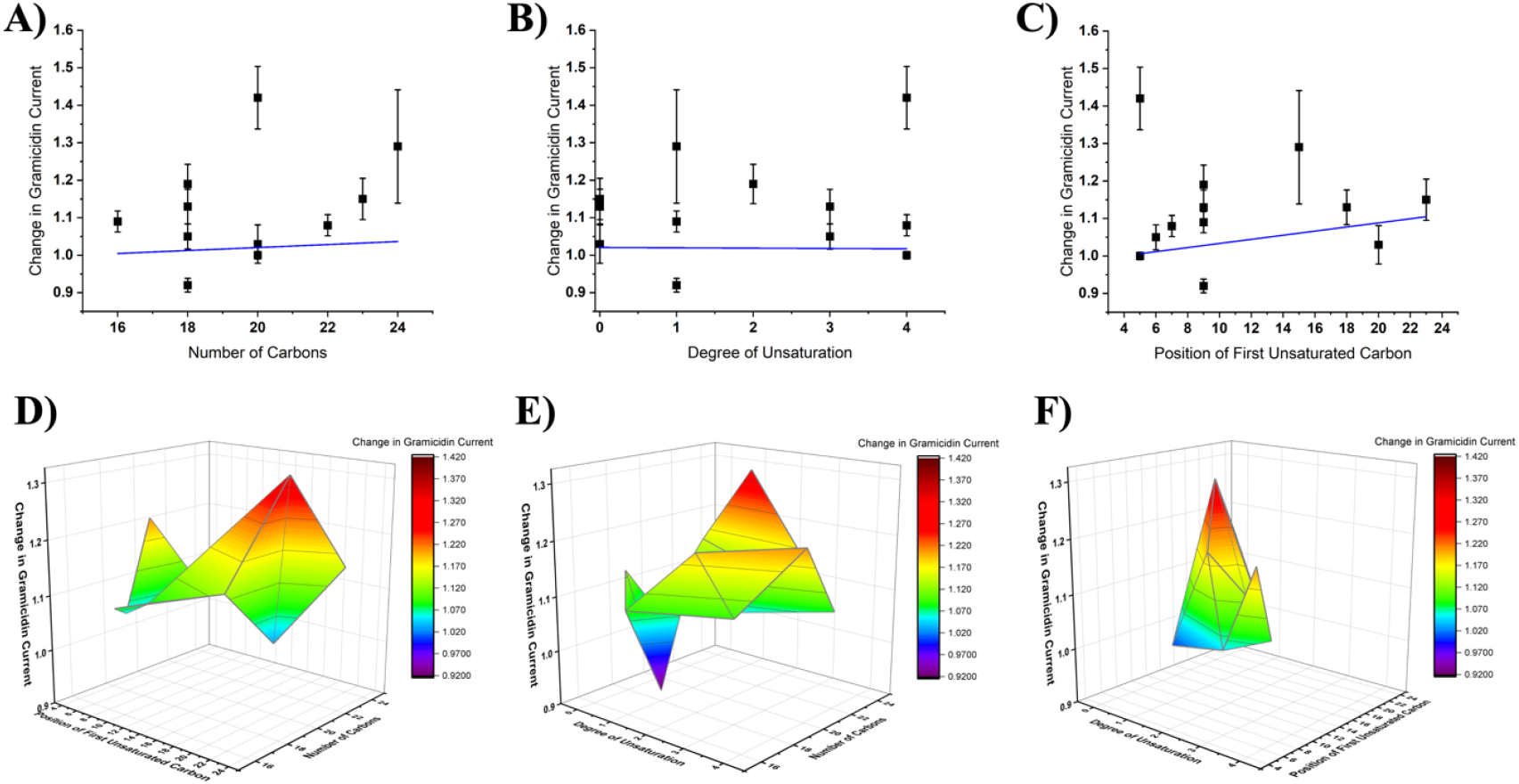
2-D linear correlation and 3D Color Surface Maps of effect of gramicidin currents vs endocannabinoid structural characteristics (FAEs). Change in gramicidin current at +150 mV plotted versus endocannabinoid properties **A)** number of carbons **B)** degree of unsaturation and **C)** the position of the first unsaturated carbon. The changes in gramicidin current did not correlate with any of these properties (P > 0.05). 3D surface plots of change in gramicidin current at +150 mV vs **D)** position of first unsaturated bond and carbon tail length **E)** number of unsaturated bonds and carbon tail length **F)** number of unsaturated bonds and position of first unsaturated bond. Minimum: OEA, 0.92, C9 position of degree of saturation, 1 degree of unsaturation and 18-carbon chain length. Maximum: oxy-AEA: 1.42. C5 position of degree of saturation, 4 degrees of unsaturation and 20-carbon chain length.

As we did for FAEs, we analyzed the relationship between the effect of 2-MG endocannabinoids on gramicidin currents and the structural properties of the lipids; chain length, degree of unsaturation, and the position of the first unsaturated bond (Fig. 5A-C). Gramicidin dimerization had a strong correlation (R: 0.92; R^2^: 0.84) with the position of the first unsaturated bond, and a moderate inverse correlation (R: -0.73; R^2^: 0.53) with the degree of unsaturation (Figure. 5 B & C, Table S5). Notably, we found that the relationship between the change in gramicidin current at +150 mV and the number of carbons in each endocannabinoid tail could be fit with a linear function (R : 0.82; R^2^: 0.67; P < 0.001). We also fit this data with a non-linear Gaussian relationship (Fig. 5A, grey), which resulted in a reduced-λ^2^ of 4.4, R^2^ of 0.73 and P = 0.07, suggesting this model does not quite reflect the relationship and a linear relationship is more appropriate. 3-dimensional surface plots reinforce this landscape in which longer tails with fewer unsaturated bonds away from the headgroup leads to maximal increase in gA currents, while shorter tails with few unsaturated bonds leads to a decrease in gA currents.

**Figure 5.**
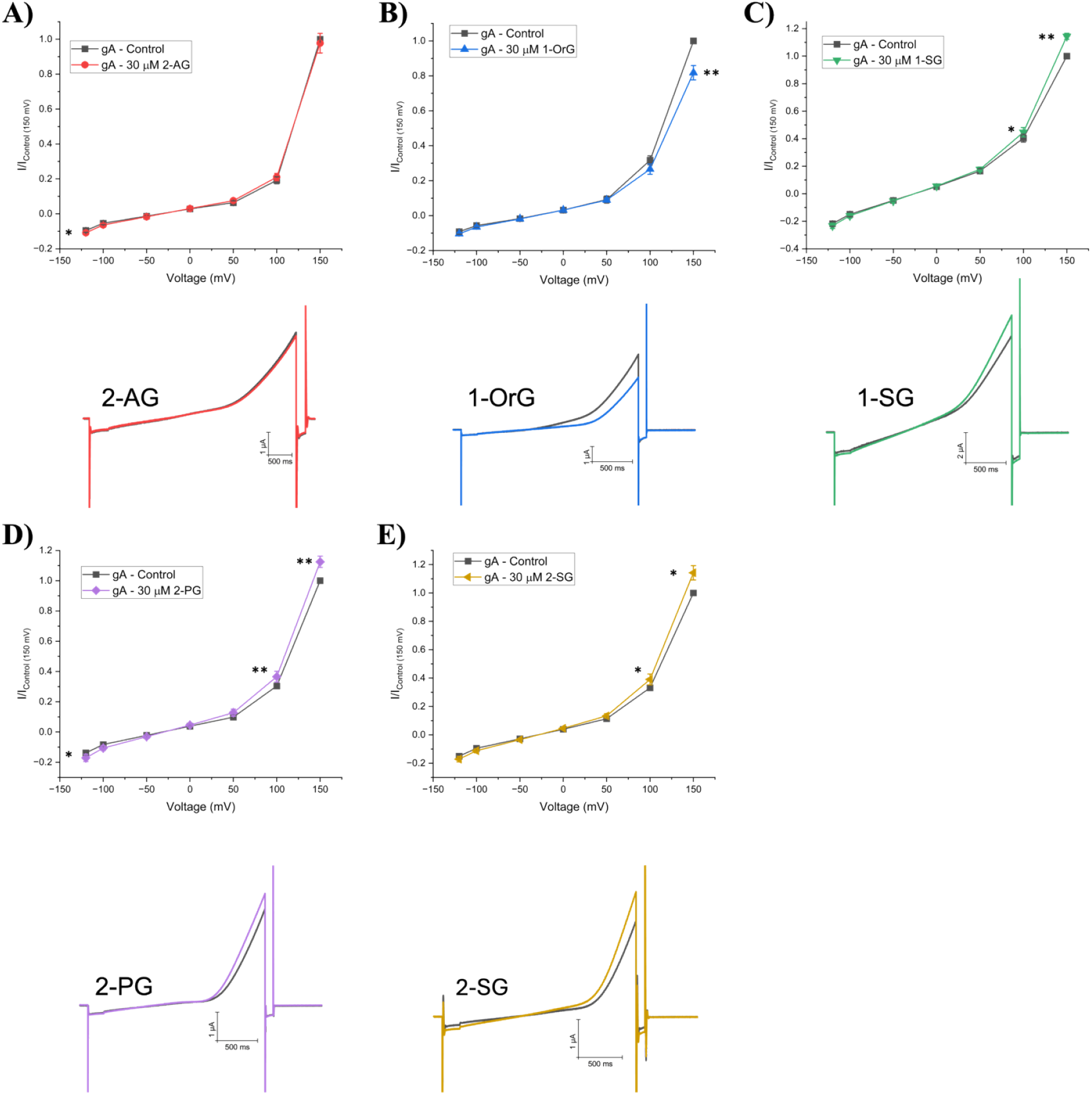
2-monoacylglycerols (2-MGs) effect on gramicidin currents (gA). (**A-E**) (*top*) Averaged cationic gramicidin currents before and after treatment with 2-AG, 1-OrG, 1-SG, 2-PG or 2-SG respectively. Test currents were averaged at -120 mV, -100 mV, -50 mV, 0 mV, +50 mV, +100 mV, and +150 mV. (*below*) Representative gA current traces before and after treatment with 2-AG, 1-OrG, 1-SG, 2-PG or 2-SG. Paired student t-test was used to determine I/I_max_ differences at -120, +100 and +150 mV (6 < n < 12; * = P < 0.05, ** = P < 0.01, *** = P < 0.001)

**Figure 6.**
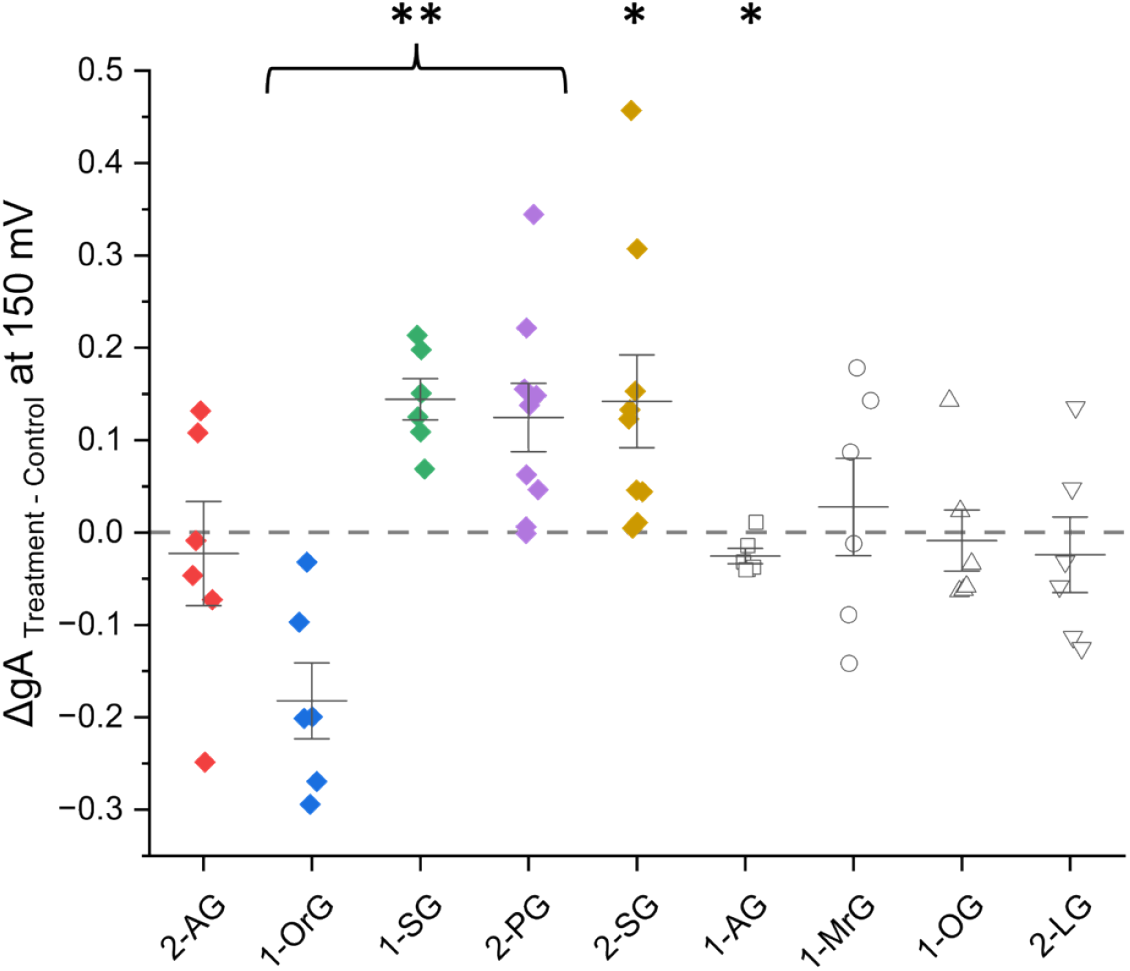
Effect of 2-monoacylglycerols (2-MGs) on gramicidin currents (gA) at +150 mV. Difference in normalized cationic currents (endocannabinoid treatment vs control). All differences are plotted for +150 mV. Paired student t-test was used to determine gA differences between treatment and control (6 < n < 12; * = P < 0.05, ** = P < 0.01, *** = P < 0.001)

**Figure 7.**
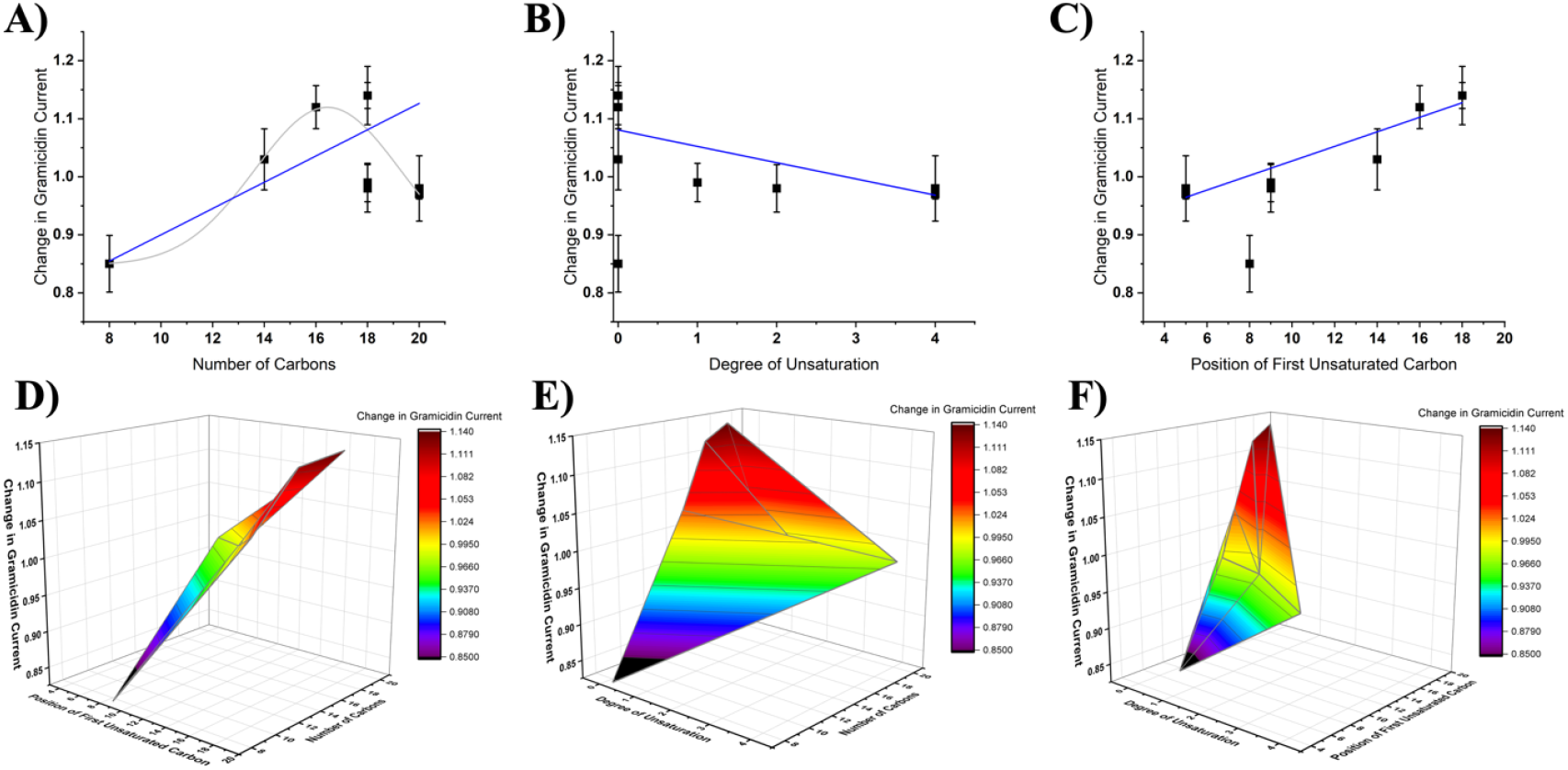
2-D linear correlation and 3D Color Surface Maps of effect of gramicidin currents vs endocannabinoid structural characteristics (2-MGs). Change in gramicidin at +150 mV current plotted versus endocannabinoid properties number of carbons **A)** degree of unsaturation **B)** and the position of the first unsaturated carbon **C)**. Change in gramicidin current had a linear correlation with the degree of unsaturation (P < 0.05) and the position of the first unsaturated bond (P < 0.001). Change in gramicidin current also linearly correlated with the number of carbons (P < 0.001) and less well with a non-linear gaussian fit (grey) (λ^2^ = 4.4). 3D surface plots of change in gramicidin current at +150 mV vs **D)** position of first unsaturated bond and carbon tail length **E)** number of unsaturated bonds and carbon tail length **F)** number of unsaturated bonds and position of first unsaturated bond. Minimum: 1-OrG, 0.82, C8 position of degree of saturation, 0 degrees of unsaturation and 8-carbon chain length. Maximum: 1-SG: 1.14. C18 position of degree of saturation, 0 degrees of unsaturation and 18-carbon chain length.

## DISCUSSION

In this study, we sought to correlate changes in membrane properties to changes in gA macroscopic currents elicited in *Xenopus laevis* oocytes. This cell model lacks CBR receptors (Karimi et al. 2018; Peshkin et al. 2019) and provides a biological lipid membrane in which we examine the effects of FAE and 2-MG endocannabinoids. Several studies have shown that gramicidin is a useful tool in determining the effects of many membrane properties such as bilayer stiffness, elasticity, thickness and deformation energy and curvature (Lundbæk et al. 2009). Some of these properties are directly correlated to the susceptibility gramicidin monomers which dimerize (to form a transmembrane cationic channel) and the current theses dimers produce. As reference amphiphilic molecules such as TX-100 (Sawyer, Koeppe, and Andersen 1989), capsaicin (Lundbaek et al. 2005) and β-OG (Sawyer, Koeppe, and Andersen 1989) tend to decrease membrane stiffness and increase elasticity. Alternatively, lipids can have varying effects for instance, cholesterol increases membrane stiffness (Lundbaek et al. 1996) and lysophosphatidylcholine (LPC) decreases membrane stiffness (Lundbaek and Andersen 1994). Notably, cholesterol is one of the only molecular entities which can modulate (decrease) membrane thickness (Lundbaek et al. 1996). Our data reveals TX-100 decreases overall cationic gramicidin current at +150 mV and +100 mV (Fig. S1 and Table S1) which agrees with previously reported data (Sawyer, Koeppe, and Andersen 1989; Lundbaek et al. 1996). Additionally, a recent study using the same experimental approach in HEK293 cells revealed TX-100 decreases overall gramicidin cationic currents while CBD increased currents (Ghovanloo, Goodchild, and Ruben 2022). Therefore, to interpret our data, we suggest that endocannabinoids which increase gramicidin current may increase membrane stiffness (decrease elasticity) while those that decrease gramicidin current similarly to TX-100 may decrease membrane stiffness (increase elasticity). Previous studies in our lab using phytocannabinoids and endocannabinoids found that 30 μM of these compounds typically elicits effects on channels expressed in *Xenopus laevis* oocytes (Mayar et al. 2024; Mayar et al. 2022). Therefore, to examine the effect of endocannabinoids on membrane properties using changes in gA current as a readout, we tested each endocannabinoid at 30 μM in this study.

Endocannabinoids may exert their effects through activating cannabinoid receptor signalling, directly binding to transmembrane proteins such as ion channels or through changes in the properties of membrane lipid bilayers. These molecules are lipophilic and like to insert themselves into the lipid bilayer to exert their effects (Oz, Yang, and Mahgoub 2022). Exogenous cannabinoids such as CBD and THC have been shown to modulate membrane elasticity (Ghovanloo et al. 2021; Ghovanloo, Goodchild, and Ruben 2022; James et al. 2022), while endocannabinoids such as anandamide (AEA) and 2-AG have also been speculated to exert their effects through changes in the lipid bilayer (Medeiros et al. 2017). Additionally, endocannabinoids and endocannabinoid-like molecules such as POEA, OEA, SEA, LEA, and 2-LG remain understudied as they have a low affinity for cannabinoid receptors CBRs (Rahman et al. 2021), and thus cannot alter intracellular signalling via that mechanism.

Fatty acid ethanolamides (FAEs) are derived from phosphatidylethanolamine (PE) which is found in the lipid membrane and contain an ethanolamide head group (Ruiz de Azua and Lutz 2019). Our data demonstrated that AEA did not alter gA currents in native oocyte membranes. Previous reports indicated that in ideal membranes, AEA increases gA single channel open lifetime and the frequency of channel appearance in planar lipid membranes (Medeiros et al. 2017) suggesting that AEA increases gramicidin dimerization and decreases membrane elasticity (increases stiffness). Thus, while AEA may be able to alter elasticity/stiffness in ideal membranes, it seems that native membranes may be less amenable to such changes. Oxy-AEA, a precursor to AEA, had the largest increase in gramicidin current followed by NEA and LEA, then smaller changes induced by DEA, POEA, SEA, TEA. The extra oxygen atom within the amide group has been speculated to increase resistance to enzymatic break down (Appendino et al. 2006). These data suggest that most FAE endocannabinoids tested increase gramicidin current may increase membrane stiffness (decrease elasticity) while OEA was the only endocannabinoid that decreased gA current and thus decreases membrane stiffness (increases elasticity). We saw no correlation between changes in gA currents and physiochemical properties of endocannabinoids; namely carbon tail length, degrees of unsaturation, and position of first unsaturated bond (Fig 3A-C). This provides further support that, FAEs likely do not exert their effects through modulating hydrophobic membrane thickness, since this type of change would likely be correlated with chain length. Rather, we anticipate changes in gA function are reflecting changes in the elastic properties of the biological membranes they are embedded.

2-monoacylglycerols (2-MGs) are derived from phosphatidylinositol 4,5-bisphosphate (PIP_2_) also found in the lipid bilayer, however they have a glycerol head group that can bind in the *sn-2* or *sn-1* positions (Ruiz de Azua and Lutz 2019; Röhrig et al. 2019). 2-PG, 1-SG, and 2-SG induce the largest increases in gA current (∼15%). These lipids have chain lengths of 16 or 18 carbons and no degrees of unsaturation. With the increase in gramicidin current we speculate 1-SnG, and 2-PG increase membrane stiffness (decrease elasticity). Interesting, 2-PG has been shown to not bind CBRs (Hanus and Mechoulam 2010) and thus could regulate the function of various membrane proteins through direct binding or changes in membrane properties. 2-AG was previously shown to increase open channel lifetime and the frequency of gA channel appearance in planar lipid membranes, however little is known about changes to cationic current (Medeiros et al. 2017). Notably, this effect of 2-AG in planar lipid bilayers was smaller than the effect observed for AEA. Our data indicates that 2-AG does not significantly modulate macroscopic gA current in the biological membranes of *Xenopus* oocytes. The lack of response from 2-AG and AEA suggest that biological membranes may be stiffer than ideal membranes, and are likely less responsive to modulators. 1-OrG, which has a short carbon tail length (C8) and no degrees of unsaturation, is the only 2-MG endocannabinoid that decreases gramicidin current.

As their name implies, 2-MGs contain a glycerol head group. Glycerol has been previously demonstrated to stiffen membranes (decreases elasticity) (Pocivavsek et al. 2011), consistent with our observation that several 2-MGs increase gA currents. We demonstrate that there are reasonable correlations between gA current changes and the number of carbons, degree of unsaturation, and the position of the first unsaturated bond of 2-MG endocannabinoids (Fig. 5A-C, Table S5). In addition to linear plots of gA changes vs. physiochemical properties of the endocannabinoids, three-dimensional plots (Figs. 3D-F and 5D-F) provide a landscape for which to better understand the contributes to membrane elastic properties. These profiles indicate that for both FAE and 2-MG endocannabinoids with longer tails with fewer unsaturated bonds that are far away from the headgroup increase membrane stiffness (or decrease membrane elasticity). However, the landscape between the 2 classes are notably different, with FAEs having more saddled features, while the landscape of 2-MGs is more smooth. Our findings indicate a potential mechanism by which endocannabinoids can exert their effects on ion channels through changes in membrane properties. More specifically, these profiles provide a backdrop to compare the landscape of changes to any ion channel’s function versus endocannabinoid properties as a mechanism to determine the contribution of changes in membrane elasticity to endocannabinoid regulation of that protein.

## ABBREVIATIONS

FAEs: fatty acid ethanolamides
AEA: Arachidonoyl Ethanolamide or anandamide
α-LnEA: α-Linolenoyl Ethanolamide
oxy-AEA: oxy-Arachidonoyl Ethanolamide
NEA: Nervonoyl Ethanolamide
OEA: Oleoyl Ethanolamide
TEA: Tricosanoyl Ethanolamide
POEA: Palmitoleoyl Ethanolamide
LEA: Linoleoyl Ethanolamide
γ-LEA: γ-Linolenoyl Ethanolamide
DEA: Docosatetraenoyl Ethanolamide
ArEA: Arachidoyl Ethanolamide
SEA: Stearoyl Ethanolamide
2-MGs: 2-monoacylglycerols
2-AG: 2-Arachidonoyl Glycerol
1-OrG: 1-Octanoyl-rac-Glycerol
1-SG: 1-Stearoyl-sn-Glycerol
2-PG: 2-Palmitoyl Glycerol
1-AG: 1-Arachidonoyl Glycerol
2-LG: 2-Linoleoyl Glycerol
2-SG: 2-Stearoyl-rac-Glycerol
1-MrG: 1-Myristoyl-rac-Glycerol
1-OG: 1-Oleoyl Glycerol
TX-100: Triton-X 100
CBD: Cannabidiol
TEVC: Two-Electrode Voltage Clamp.

